# Activation of IL1A/IRAK1 axis and downstream proinflammatory signaling in healthy adult and neonatal African American skin

**DOI:** 10.1101/2024.11.18.624144

**Authors:** Dimitri Trubetskoy, Patrick Grudzien, Anna Klopot, Lam C Tsoi, Roopal V Kundu, Bethany E Perez White, Irina Budunova

## Abstract

Differences in prevalence of inflammatory skin diseases including atopic dermatitis and psoriasis in African American (AA) versus White Non-Hispanic (WNH) population are well recognized. However, the underlying mechanisms are largely unknown. We previously observed significant differences in healthy AA skin transcriptome with differentially expressed genes (DEG) enriched for inflammation and cornification processes. Here we analyzed proteome in skin biopsies from healthy AA and WNH volunteers using *Olink*® *Explore Inflammation* 384 biomarker panel. Among proteins with higher expression in AA skin were IRAK1, IL1A, IL4, IL22RA1. IL1A binding to IL1R1 receptor is known to result in recruitment of adapter molecules such as IRAK1, and activation of downstream NF-κB and MAPK signaling. We confirmed NF-κB and ERK1/2 activation in AA skin by Western blot analysis of their phosphorylation at specific activating sites. Importantly, we observed similar differences between AA and WNH neonatal foreskin and between AA and WNH 3D skin organoids. Further analysis of DEG promoters by Gene Transcription Regulation Database (GTRD) pointed to NF-κB and AP1 as key transcription factors involved in AA DEG regulation. Overall, proinflammatory signaling in healthy AA skin starting early in childhood may contribute to the increased risk of certain inflammatory skin diseases within the AA population.

## INTRODUCTION

Morphological and physiological differences in African American (AA) skin compared to White non-Hispanic (WNH) skin are well-documented. AA skin is characterized by accumulation of eumelanin pigment, numerous convoluted dermal-epidermal junctions, increased epidermal proliferation, thicker stratum corneum, stronger corneocyte cohesion, and enhanced skin barrier function (Brown-Korsah et al., 2022; Halder & Nootheti, 2003; Salminen et al., 2023).

There are also differences in the prevalence and clinical manifestations of many inflammatory skin diseases, abnormal wound healing and skin cancer. The risk of atopic dermatitis (AD), acne, hidradenitis suppurativa, and lupus erythematosus as well as keloids (abnormal scarring) is significantly higher in AA populations (Agbai et al., 2014; Alexis & Blackcloud, 2014; Eichenfield et al., 2014; Feldman et al., 2019; Halder & Nootheti, 2003; Mei-Yen Yong & Tay, 2017; Price et al., 2020). WNH individuals have a higher risk of psoriasis, however, the severity of psoriasis might be greater in AA patients (Abrouk et al., 2017; Yan et al., 2018). This disparity could be even more pronounced, given the challenges in visualizing and scoring erythema in darker skin (Adawi et al., 2023; Kaufman & Alexis, 2018).

The most extensive studies of molecular phenotype of AA diseased skin were done in AA patients with AD. Transcriptomic analysis of full-thickness skin biopsies or tape strips revealed numerous differentially expressed genes (DEG) in AA lesional skin compared to AA non-lesional or healthy skin (Hawkins et al., 2024; Luo et al., 2007; Navrazhina et al., 2024; Nomura et al., 2020; Wu et al., 2020). In some of these studies RNA-seq was combined with analysis of serum and skin immunostaining, and the results were compared with WNH AD skin phenotype (Brown-Korsah et al., 2022; Brunner & Guttman-Yassky, 2019; Wongvibulsin et al., 2021). These studies revealed that AA AD patients often exhibit higher serum IgE levels, higher CRP (C-reactive protein) values, ferritin, and blood eosinophils compared to WNH AD patients along with immune polarization primarily characterized by T_H_2/T_H_22 skewing. In addition, it was found that filaggrin (FLG) loss-of-function mutations typical for WNH and Asian AD patients are not prevalent in AA patients. Instead, there are frequent loss-of-function filaggrin 2 (FLG2) mutations and significantly lower levels of ceramides in the stratum corneum in AA patients with AD (Brown-Korsah et al., 2022; Brunner & Guttman-Yassky, 2019). Interestingly, the skin phenotype of Tanzanian AD patients appeared to be consistent with that of AA patients, exhibiting dominant T_H_2/T_H_22 skewing, minimal dysregulation of terminal differentiation with no suppression of epidermal barrier marker genes, such as FLG, loricrin, and periplakin but with significant down-regulation of genes related to lipid metabolism including HAO2, ELOVL3, GAL, FAR2, AWAT1 (Lang et al., 2021)

However, despite well-recognized morphological and physiological differences, the molecular landscape of healthy AA skin compared to WNH or other skin types remains underexplored. There is practically no data on healthy AA skin proteome and very limited literature on genomic profiling comparing healthy AA skin to WNH skin, except for some transcriptomic studies with a minimal number of volunteers/group whose skin biopsies were mostly used as a reference for comparison with the same ethnicity diseased skin features/expression status (Sanyal et al., 2019; Walker et al., 2020; Wongvibulsin et al., 2021).

In our recent study, the analysis of gene expression in skin biopsies from healthy AA and WNH volunteers of both sexes using RNA-sequencing identified hundreds of differentially expressed genes (DEGs, FC>1.5; P<0.02) in AA skin (Klopot et al., 2022). Among those, we observed upregulation of FCER1G - the major IgE receptor; proinflammatory genes such as TNFα and IL-32; and epidermal differentiation cluster (EDC) genes like late cornified envelope (LCE) genes. Notably, the expression of some of these DEGs could be correlated with clinical observations, such as increased IgG and IgE levels in the serum of healthy AA individuals and AA patients with various diseases (Albandar et al., 2002; Lucey et al., 1992; Rinker et al., 2007). Additionally, the elevated expression of EDC genes could be linked to the increased stratum corneum thickness and better barrier function in AA skin (Girardeau-Hubert et al., 2019; Muizzuddin et al., 2010). Notably, only four known pigment-related genes were found among the DEGs in AA skin (Crawford et al., 2017), indicating that differential gene expression in the skin is not closely associated with melanogenesis.

The goals of our study were to (i) confirm previously found in our studies overexpression of selected proinflammatory genes in skin biopsies from a different group of volunteers; (ii) expand upon these findings by analyzing protein expression in AA and WNH healthy skin using Olink™ proteomics (Uppsala, Sweden) and Western blotting in full-thickness adult skin biopsies; (iii) identify upstream pathways driving proinflammatory signaling in healthy AA skin. We also included neonatal skin in our studies, that differs from adult skin in gene/protein expression and barrier function (Visscher et al., 2022; Visscher et al., 2021) to assess whether intrinsic proinflammatory signaling in AA skin develops earlier in life.

We revealed the increased expression of numerous proinflammatory proteins in AA skin, activation of the IL1A/IRAK1 axis and downstream inflammatory signaling pathways mediated by NF-κB and MAPKs in both adult and neonatal skin, and confirmed overexpression of TNFα, FCER1G (receptor for IgE), IL1A and LCE genes in both types of AA skin.

## RESULTS

### Olink Proteomics analysis of inflammation-related protein expression in adult skin

We extended our previous findings on intrinsic proinflammatory signaling in AA skin by Olink^TM^ proteomics (Uppsala, Sweden) in additional full-thickness skin biopsies from 12 healthy volunteers (5 AA and 7 WNH women, aged 24-41 years) grouped by self-identification combined with Fitzpatrick skin score. Published data indicate that self-reported African American ancestry correlates with genotyping at approximately 95% (Yaeger et al., 2008).

The analysis using the Explore inflammation 384 biomarker panel confirmed the existence of proinflammatory signaling in AA skin and revealed 11 differentially expressed proteins (DEP, FC >1.5; P<0.05). Nine out of these DEPs were upregulated in AA skin including known mediators of skin inflammation IL4, IL1A, IL22RA1 (a component of the receptor for IL20, 22 and 24) and IRAK1 whose expression was 2-4 folds higher in AA skin (**Fig. 1 A**).

**Figure 1.**
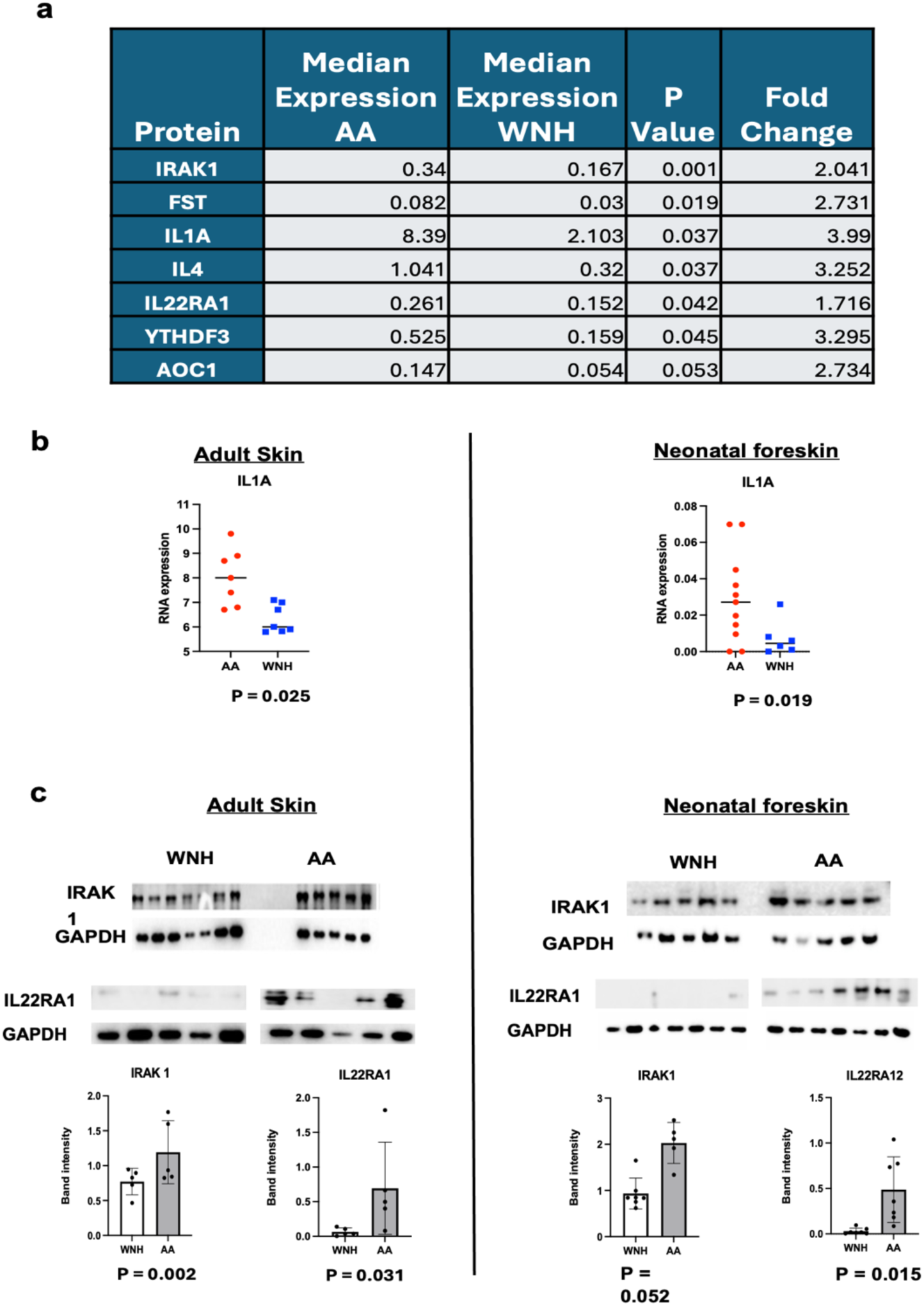
Activation of proinflammatory IL1A/IRAK1 axis in AA skin. Proteins and RNA were extracted from 4 mm full-thickness skin biopsies from the upper inner arm skin in AA and WNH healthy volunteers and from neonatal foreskin. Protein extracts were used for Olink proteomics (**A**). and Western blotting (**C**). GAPDH was used as a loading control. **B.** IL1A expression was assessed by qRT-PCR in individual AA (red circles) and WNH (blue squares) adult skin RNA samples. RPL27 was used as a cDNA normalization control. Statistical analysis was performed using unpaired two-tailed t test.

Several other DEPs (Suppl Table 1) whose function in skin/skin inflammation has been less studied included gonadal protein, Follistatin (FST), an inhibitor of follicle-stimulating hormone release. It was shown that FST is highly expressed in keratinocytes and skin fibroblasts, where it negatively regulates the signaling of TGF-β family members activin and bone morphogenic proteins (BMP) and is involved in skin and appendages development, wound healing and inflammation (Antsiferova et al., 2009). Another important DEP is Copper-containing Diamine Oxidase 1 (AOC1/DAO) - a membrane glycoprotein that deaminates histamine and related molecules and is involved in allergic reactions in skin and other organs (Comas-Baste et al., 2020; Okutan et al., 2023). RABGAP1L (RAB GTPase Activating Protein) and RNA binding protein YTHDF3 have been linked to the development of skin cancer (Shi et al., 2022). Unexpectedly, the expression of NELL2 protein, a marker for AD (Yu & Li, 2022) was lower in AA healthy skin.

### Activation of IL1R1/IRAK1 axis and downstream NF-κB and MAPK signaling in adult and neonatal AA skin

Binding of IL1A to IL1R1 receptor results in the recruitment of adapter molecules such as IRAK1, IRAK4, MYD88, and activation of downstream NF-κB and MAPK signaling cascades (Cohen et al., 2015; Di Paolo & Shayakhmetov, 2016; Jain et al., 2014; Malik & Kanneganti, 2018). Thus, we first focused on validating IL1A and IRAK1 overexpression in AA skin. We added neonatal foreskin to our studies (7 AA and 7 WNH samples, grouped by mothers’ self-identification) to include male skin for the analysis to assess whether proinflammatory signaling in AA skin is developed earlier in life when skin has different features, not completely established barrier function and a markedly dissimilar array of protein biomarkers (Visscher et al., 2021).

Even though the differences in morphology of adult AA and WNH skin are well-documented (Girardeau et al., 2009; Montagna & Carlisle, 1991), the comparative analysis of neonatal AA and WNH skin is lacking. Thus, we first performed morphological analysis of AA and WNH neonatal foreskin focusing on rete ridges (convoluted dermal-epidermal junctions), total thickness of epidermis, thickness of stratum corneum (cornified layer). As shown in Fig. **2**, the features of neonatal AA skin were very similar to those described in AA adult skin and included statistically significant increased frequency and depth of rete ridges and increased stratum corneum thickness compared to WNH neonatal skin. In contrast to the reports that keratinocyte proliferation in adult AA skin is increased (Girardeau-Hubert et al., 2019), Western blot analysis of cell cycle marker PCNA did not reveal the increased proliferation in neonatal AA skin (Fig. 2C).

**Figure 2.**
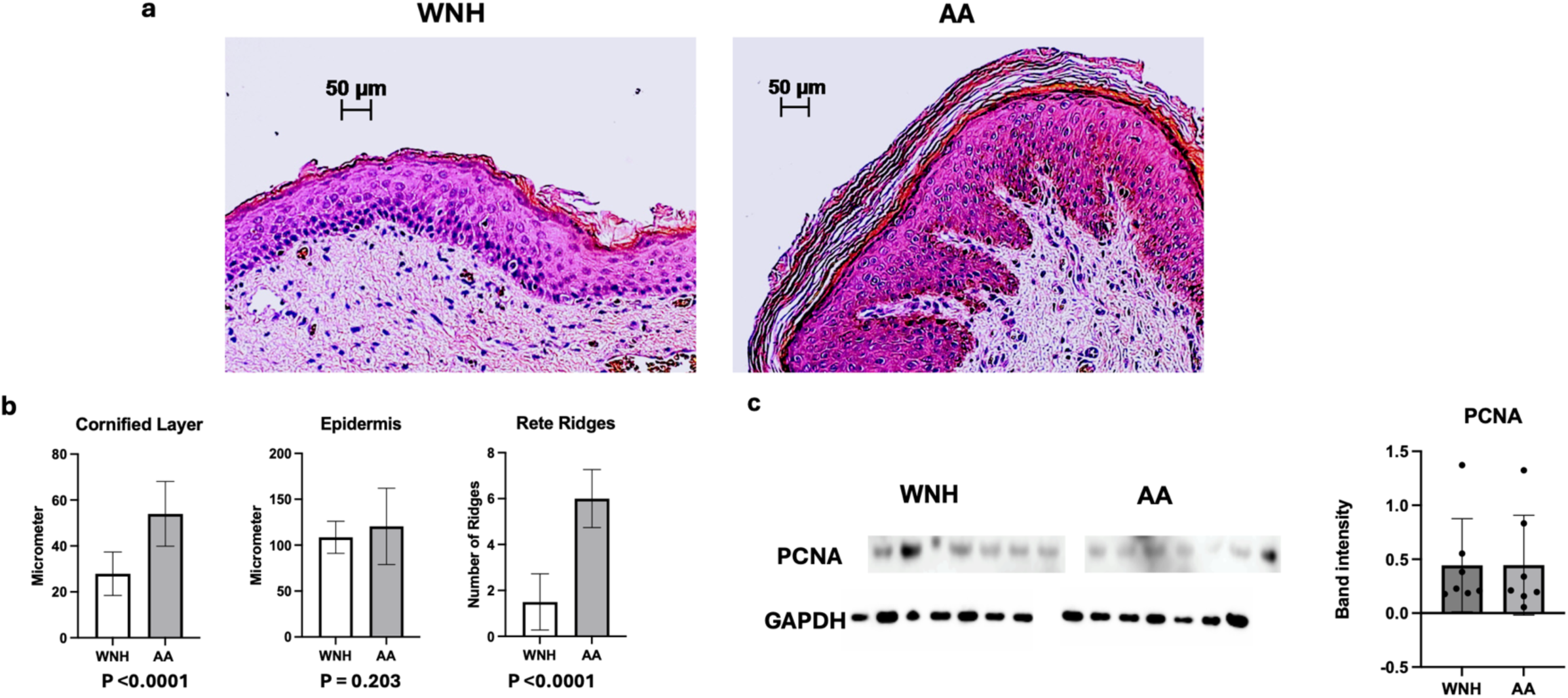
Differences in AA and WNH neonatal skin morphology. **A**. Hematoxylin and eosin (H&E) staining was done in formalin-fixed, paraffin-embedded samples of neonatal skin. **B**. Total epidermal thickness, thickness of cornified layer and number of rete ridges/field of view were determined by ZeissAxio Vision software. At least three individual fields of view/slide in 7 AA and 7 WNH individual skin samples (∼ 20 images/group) were analyzed. Statistical analysis. **C.** Western blot analysis of proliferative marker PCNA expression in AA and WNH neonatal skin protein extracts. Statistical analysis was performed using unpaired two-tailed t test.

Western blotting revealed the increased expression of IRAK1 and IL22RA1 proteins in AA adult and neonatal skin (**Fig**. **1C**), The expression of IL1A and IL4 were below Western Blot detection, however, IL1A expression at mRNA was significantly increased in both adult and neonatal AA skin (**Fig. 1B**).

Next, we assessed the activation of downstream NFκB and MAPK signaling. NFκB activation is a complex multi-step process. The primary mechanism for canonical NFκB activation is the inducible degradation of IκBα triggered through its site-specific phosphorylation (at Ser32 and Ser36) by a multi-subunit IκB kinase (IKK) complex composed of two catalytic subunits, IKKα and IKKβ. In addition, IKKβ phosphorylates major NFκB protein RelA/p65 at Ser536 and Ser468 which facilitate RelA/p65 nuclear translocation and enhance NFκB transcriptional activity (Christian et al., 2016; Motolani et al., 2020).

MAPKs (the mitogen-activated protein kinases) including extracellular signal-regulated kinase (ERK1/2), Jun kinase (JNK/SAPK) and p38, are activated via phosphorylation cascade induced by extracellular stimuli (growth factors, hormones, stress) and transmitted first to MAPKKKs then to MAPKKs and finally to MAPKs via sequential phosphorylation of these kinases at specific amino acids (Seger & Krebs, 1995).

We assessed activation of IKKs, IκBα, p65/RelA and conventional MAPKs (ERK1/2, JNK1/2 and P38) kinases by Western blotting using Abs against specific activating phosphorylation sites in these proteins. We indeed observed the constitutive activation of NF-κB in healthy adult **(Fig. 3)** and neonatal **(Fig. 4**) AA skin as determined by the statistically significant increased phosphorylation of IKKα/β, RelA/p65 at Ser536 and Ser468 and IκBα at Ser32 that targets IκBα for ubiquitination and proteolysis. Analysis of major MAPK signaling branches by Western blotting with respective phospho-Abs revealed significant increase in ERK1/2 activation in both AA skin types and JNK1/2 activation in AA newborn skin (Figs 3 and 4). We were not able to detect WB signal related to p38 phosphorylation. Despite the substantial individual variability in phosphorylation patterns, the increase of phosphorylation in NFκB-related proteins was in most cases statistically significant.

**Figure 3.**
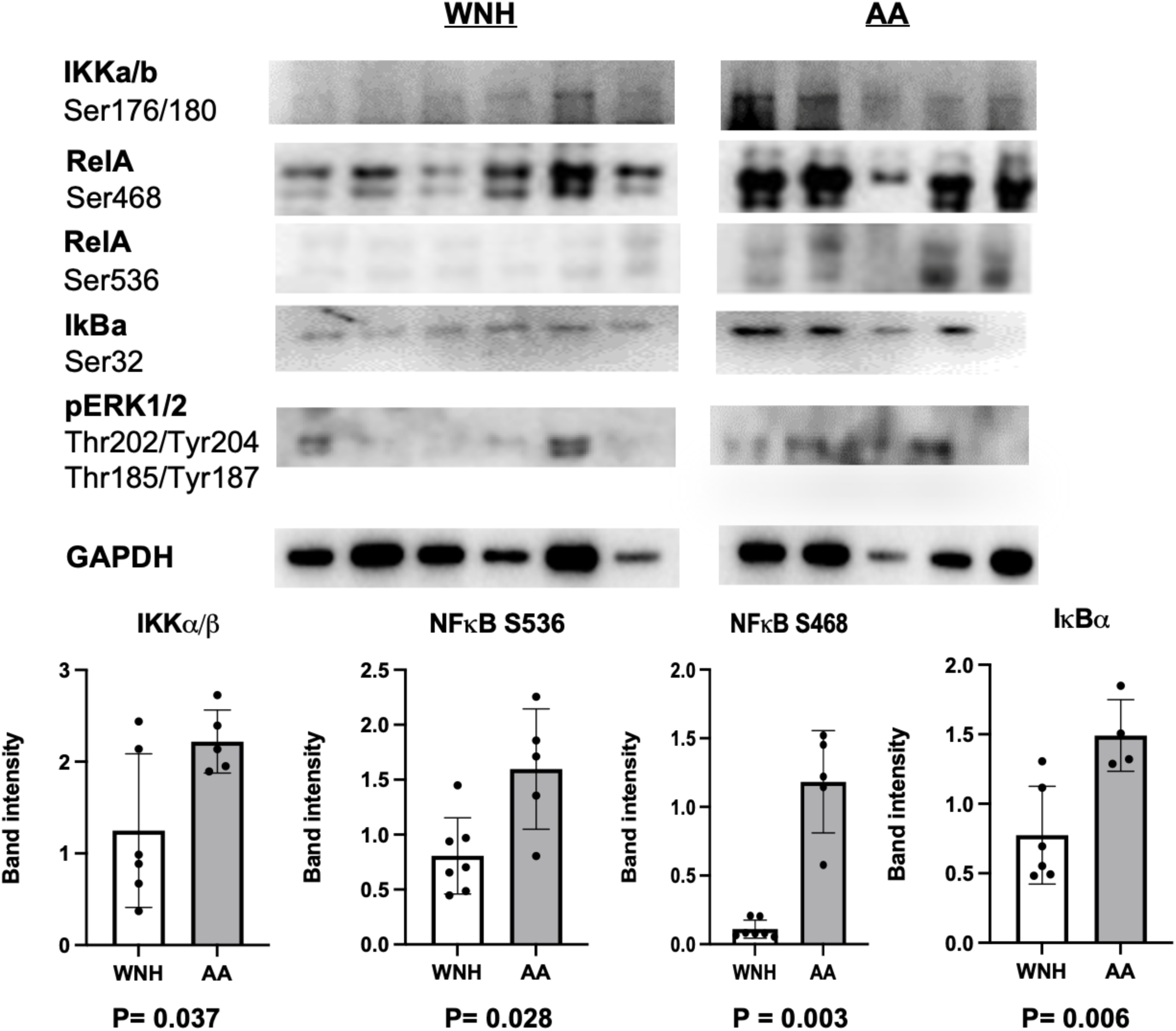
Activation of proinflammatory NF-kB and MAPK signaling pathways in adult African American skin. Proteins extracts from AA (red circles) and WNH (blue squares) adult skin biopsies were used for Western blotting. GAPDH was used as a loading control. Statistical analysis was performed using unpaired two-tailed t test.

**Figure 4.**
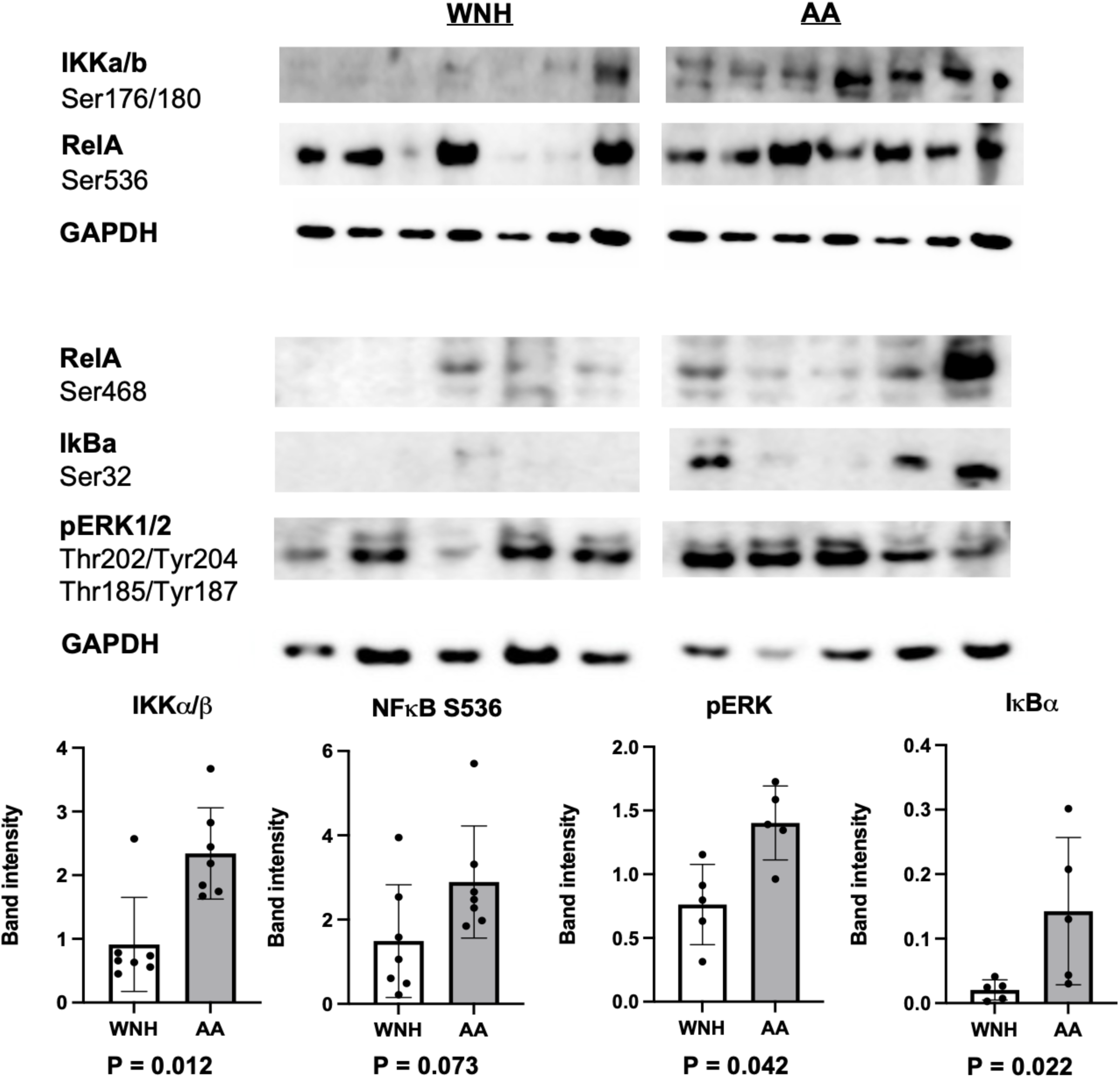
Activation of proinflammatory NF-kB and MAPK signaling pathways in African American neonatal foreskin. Proteins extracts from AA (red circles) and WNH (blue squares) neonatal foreskin were used for Western blotting. GAPDH was used as a loading control. Statistical analysis was performed using unpaired two-tailed t test.

### Analysis of IL1R1/IRAK1 axis and downstream signaling in AA and WNH human skin organoids (HSO)

The skin consists of multiple cell types. Thus, to address the role of keratinocytes in activation of IL1A/IRAK1 axis and downstream signaling in human skin we compared the expression of IRAK1, NF-κB and MAPK phosphoproteins in AA and WNH human skin organoids (HSO) made from the individual primary keratinocyte lines. The overall difference in protein expression/phosphorylation between AA and WNH HSO was less pronounced compared to full-thickness skin, and there was only minor increase in IRAK1 expression in AA HSO (**Fig. 5**). However, the increase in phosphorylation of p65/RelA, IκBα and ERK1/2 in AA HSO was statistically significant. (**Fig. 5**)

**Figure 5.**
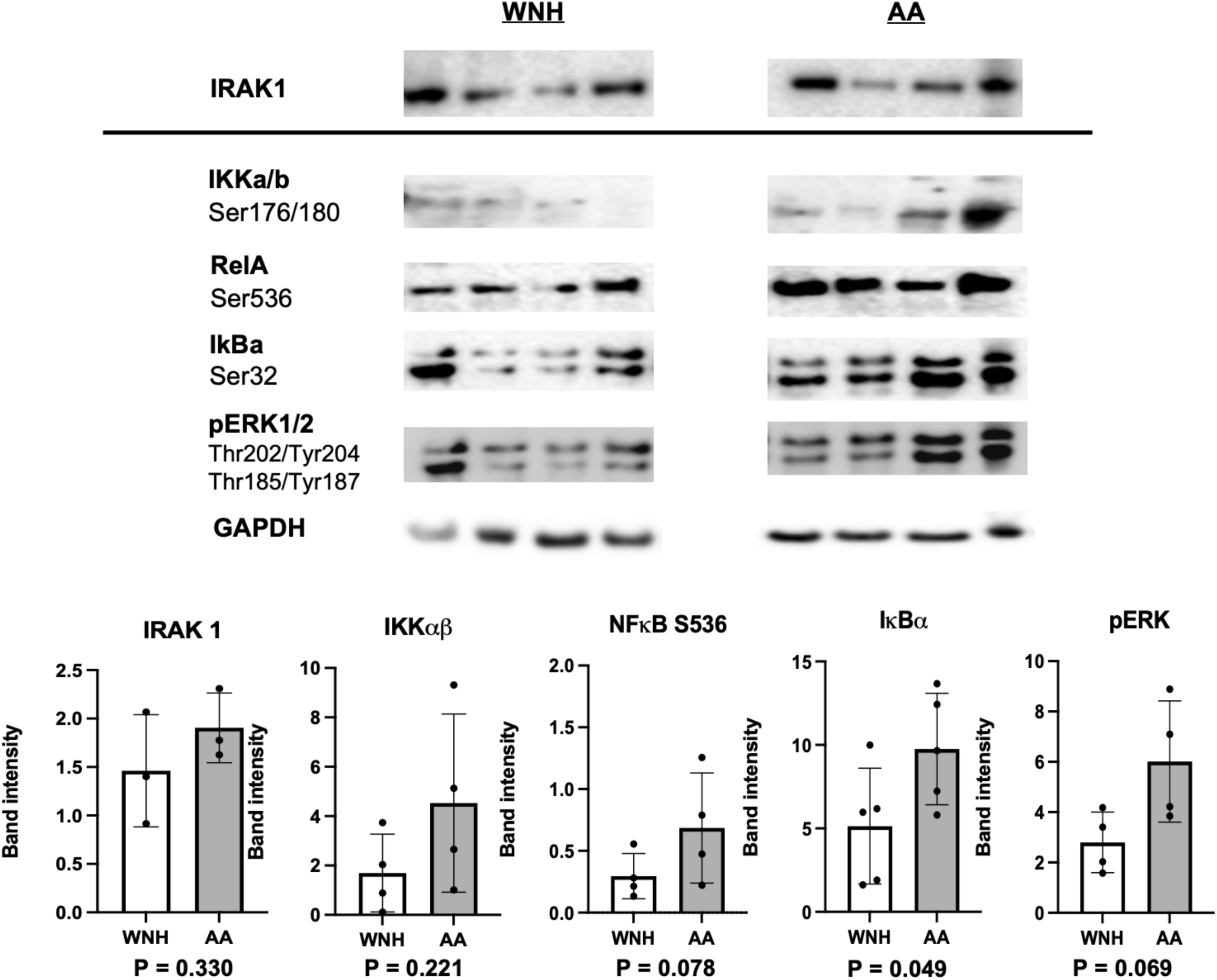
Activation of proinflammatory NF-kB and MAPK signaling pathways in African American human skin organoids (HSO). Proteins extracts from the epidermal compartment of AA and WNH HSO were used for Western blotting. GAPDH was used as a loading control. Statistical analysis was performed using unpaired two-tailed t test.

### Differential gene expression in adult and neonatal AA skin compared to WNH skin

We previously reported significant differences in AA and WNH skin transcriptome with DEGs in AA skin enriched by inflammation and formation of cornified envelope functions (Klopot et al., 2022). This previous transcriptome analysis was done in skin biopsies obtained from volunteers of both sexes aged 25-64. The skin samples for proteomics were obtained from a different, more focused group of volunteers, all females, aged 24-41 yeas. Thus, we took the opportunity to determine whether the expression difference in proinflammatory genes and keratinocyte differentiation markers such as LCEs is reproducible. Q-PCR analysis of RNA extracted from the same biopsies that were used for Olink proteomics confirmed the increased expression of TNFα, FCER1G (major Receptor for IgE), CCL2, several LCE genes (LCE1E, 2B, 5A) along with IL1A and MAPK11/p38b in AA adult skin (**Fig. 6A**). Importantly, TNFα, FCER1G and IL1A mRNAs were also highly expressed in AA neonatal skin (**Fig. 6B**). However, we did not reveal significant differences in LCE genes expression in AA versus WNH neonatal skin possibly due to incomplete maturation of skin in newborns (Fluhr et al., 2010; Rahma & Lane, 2022).

**Figure 6.**
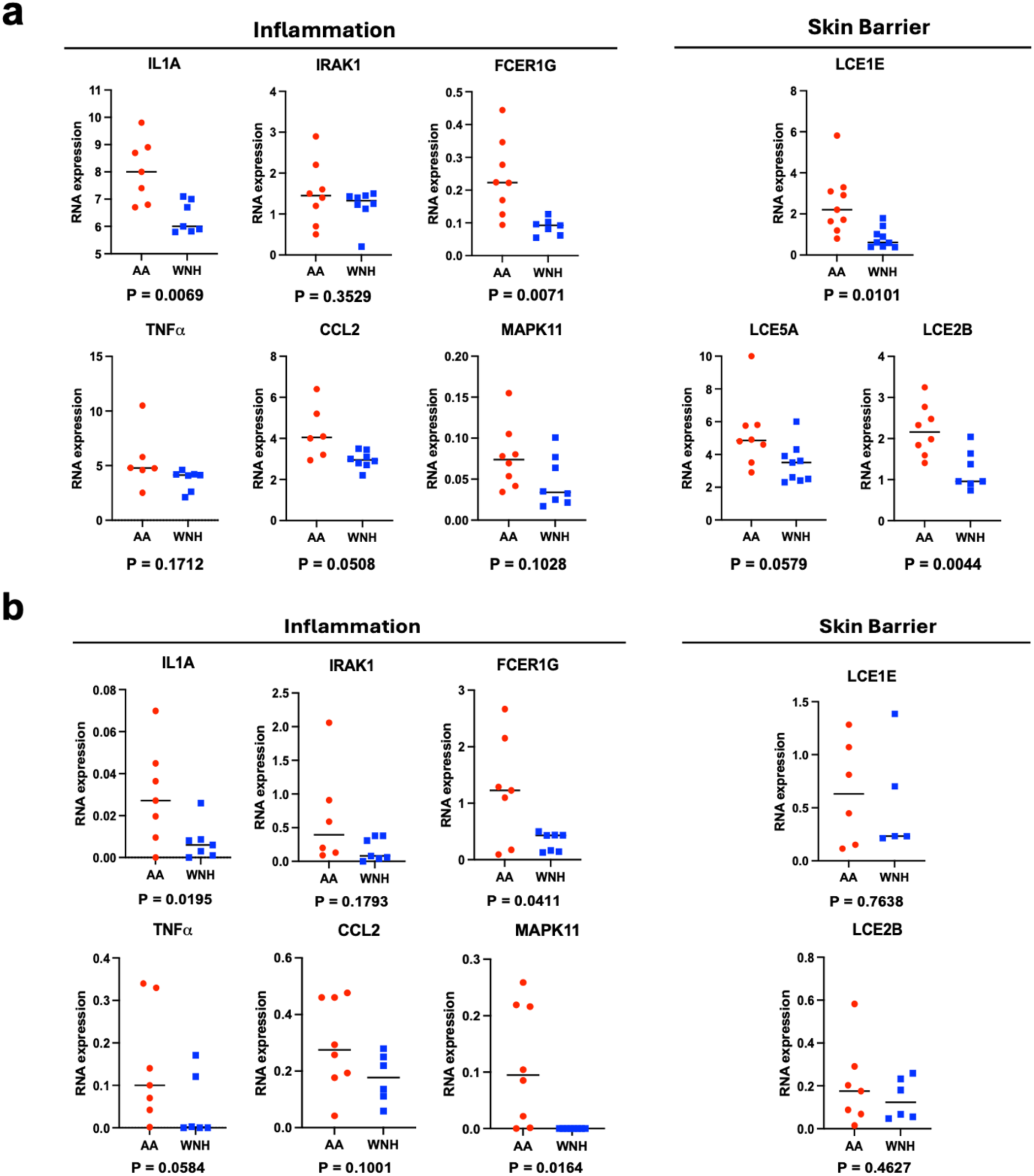
Increased inflammatory and skin barrier gene expression in AA and WNH adult and neonatal skin. RNA was extracted from full-thickness skin biopsies from the upper arm skin in AA and WNH healthy volunteers **(A**) and AA and WNH neonatal foreskin (**B**). Gene expression was assessed by qRT-PCR in individual AA (red circles) versus WNH (blue squares) adult and neonatal skin RNA samples. RPL27 was used as a cDNA normalization control. Statistical analysis was performed using unpaired two-tailed t test.

### Analysis of NFκB and AP1 putative binding sites in AA skin DEG promoters

The major transcription factors activated by signaling cascades related to IL1A/IRAK1 axis are NF-κB and AP1 (Activator Protein-1) downstream from MAPKs. We used our previously published bulk RNA-seq data (Lili et al., 2019), [GSE120783] to assess the role of these transcription factors in the regulation of AA DEGs expression. We searched for putative NF-κB (RelA) and AP1 (c-Fos and c-Jun separately) binding sites in DEG promoters using GTRD (Gene Transcription Regulation Database) -database that combines the results of ∼ 23,000 ChIP-Seq experiments performed in 30 cell types, and reports site count information for 65,000 transcribed DNA regions with ENSEMBL IDs representing coding and noncoding RNAs. For transcription factor (TF) binding sites, we selected meta-cluster analysis and a maximum gene distance of 5000 nucleotides to define transcription initiation sites in DEGs. To estimate the number of putative binding sites for the TFs RELA (P65), cFOS and cJUN in DEG promoters, we searched within (−100, 1000 bases) from the start codon. DEGs with ≥ 2 binding sites for specific TF were considered TF targets. The GTRD database is not tissue/cell type specific. About 32% of all DEGs emerged as putative RELA targets, 21% were putative cJUN and 11 % cFOS targets with at least two chromatin immunoprecipitation (ChIP)-Seq peaks in their promoters (**Suppl. Fig. 1**).

## DISCUSSION

The major goal of this work was to further explore the proinflammatory bias in healthy AA skin by proteomics. The Olink proteomics analysis of skin biopsies followed by Western blot validation provided novel, clinically important information on the healthy AA skin protein expression, revealed the activation of IL1R1/IRAK1 axis and the activation of downstream proinflammatory NF-κB and MAPKs signaling. In addition to their leading role in inflammation and cutaneous immunity, NF-κB and MAPKs together with downstream AP1 transcription factors are involved in control of skin morphogenesis, affect skin cells proliferation, survival, differentiation and regulate skin barrier function (Sur et al., 2008; Wullaert et al., 2011; Young et al., 2017). Therefore, it is plausible that some differences in the morphology and physiology of healthy AA skin result from heightened activation of NF-κB and MAPKs/AP1.

We acknowledge that our analysis of differential protein expression in adult skin has some limitations due to the relatively small size of volunteer cohort. Nevertheless, we detected a similar increase of IRAK1 expression at protein level and IL1 A at mRNA level along with the activation of NF-κB and MAPKs in neonatal AA skin suggesting that proinflammatory signaling in AA skin is established very early in life. This is a novel important finding as comparative analysis of gene/protein expression in neonatal skin of color has not been done previously. Experiments with 3D HSO cultures further confirmed this major observation.

In this work we focused only on two out of 11 DEPs proteins, IL1A and IRAK1, as they act cooperatively within the same major pro-inflammation signaling network activated by IL1R1. In addition, IL1A and IRAK have other important functions and may activate inflammatory signaling via different mechanisms. The truncated IL1A can enter the nucleus to induce the expression of proinflammatory cytokines such as IL6 and IL8 and some chemokines independently of IL1R1 (Cohen et al., 2015; Malik & Kanneganti, 2018). In turn, IRAK1, a serine/threonine-protein kinase that can phosphorylate multiple targets including E3 ubiquitin ligases, TIRAP – an adaptor protein involved in the TLR4 signaling pathway; the interferon regulatory factor 7 (IRF7) resulting in transcriptional activation of type I IFN genes and STAT3 (Kim et al., 2024).

Other important DEPs revealed by proteomics, include IL22RA1 and IL4 that could contribute to activation of proinflammatory signaling in healthy AA skin. IL22RA1 is a component of the receptor for IL22, IL20 and IL24. IL22RA1 enables IL22 to activate ERK1/2, Akt, JAK/STAT pathways. Activation of IL-22 has been associated with keratinocyte proliferation, epidermal hyperplasia, hyperkeratosis, and other barrier abnormalities and contributes to the development of AD, allergic contact dermatitis and psoriasis (Laska et al., 2024; Wongvibulsin et al., 2021). As discussed above, Th22 skewing is typical for AD in AA and in Tanzanian African patients (Lang et al., 2021).

JAK/STAT pathway plays a critical role in orchestrating immune and inflammatory responses. Over the years, the JAK/STAT pathway has been extensively investigated due to its key role in the pathogenesis of several chronic inflammatory conditions, e.g. psoriatic arthritis, AD, psoriasis, and inflammatory bowel diseases. Taking into consideration the importance of JAK/STAT signaling in the development of inflammatory skin diseases and the emerging importance of JAK/STAT inhibitors for the treatment of different immune skin diseases such as AD, alopecia areata, vitiligo, hidradenitis suppurative that have higher risk for AA population, the analysis of Th22 and JAK/STAT signaling in healthy AA skin will be an important future focus of research.

In conclusion, this work offers new insights into the molecular landscape of healthy AA skin, revealing the intrinsic proinflammatory bias and emphasizing the importance of developing targeted interventions for the precision prevention and treatment of inflammatory skin diseases prevalent in the AA population.

## MATERIALS AND METHODS

The detailed Materials and Methods are presented in the Supplement.

### Human skin samples

Full-thickness adult skin biopsies and neonatal skin samples were provided by Northwestern Skin Biology and Diseases Resource-Based Center (SBDRC) through the IRB-approved Dermatology Tissue Acquisition and Repository services. Biopsies and other skin samples were cut into several sections for protein and RNA extraction.

### 3D human skin organoids (HSO)

HSO made from neonatal human epidermal keratinocytes (NHEKs) were provided by SBDRC STEM Core. HSOs were cultured as described earlier (Simpson et al., 2010).

### Morphometric analysis of neonatal skin

Total epidermal thickness, thickness of cornified layer and number of rete ridges/field of view were analyzed by ZeissAxio Vision software in formalin-fixed paraffin-embedded neonatal skin. At least three individual fields of view/slide in seven AA and seven WNH individual skin samples were analyzed.

### Olink proteomics

Part of snap-frozen adult skin biopsies were homogenized in a Qiagen Tissue PowerLyzer at 3,500 rpm in TRIzol according to the manufacturer’s protocol (Thermo Fisher Scientific). Total protein was isolated according to manufacturer’s protocol and eluted in 1X PBS. 50µl protein extracts diluted to 1mg/ml/sample were loaded on a 96-well PCR plate and submitted to Olink for quantification of 384 proteins according to their Explore 384 Inflammation panel. Differential protein expression (DEP) analyses were performed using R software (www.R-project.org).

### Protein Extraction and Western Blotting

Proteins from the part of adult skin biopsies, neonatal foreskin and epidermal compartment of HSO were extracted with Urea Sample Buffer (USB). Protein concentration analysis was determined by Amido Black Assay.

Proteins were resolved by SDS-PAGE and transferred to nitrocellulose membranes. After blocking with TBST:5% non-fat milk (Bio-Rad) membranes were incubated overnight at 4 °C with primary antibodies overnight following incubation with secondary antibodies at room temperature. Membranes were imaged using AZURE ECL reagents and the AZURE Chemidoc digital imager and equal loading was normalized by GAPDH expression.

We used following antibodies for Western blotting: IRAK1 (Cell Sig #4504), NFκB S536 (Cell Sig #3033), NFκB S468 (Cell Sig #3039), IKKα/β (Cell Sig #2697), IkBa (Cell Sig # 2859), MAPKs (ERK 1/2, JNK, p38)(Cell Sig #4695, Cell Sig #4668, Cell Sig #4511), GAPDH (Cell Sig #2118), anti-Rabbit IgG (Cell Sig #7074P2). pERK1/2 Ab is specific for Thr202/Tyr204 in ERK1 and Thr185/Tyr187 in ERK2, pJNK antibody detects SAPK/JNK at Thr183 and Tyr185, and p-P38 antibody detects phosphorylation at Thr180/Tyr182 of any p38 isoforms (p38α, p38β, p38γ, p38δ), PCNA (Cell Sig #2586), IL22RA1 (Proteintech #13462-1-AP).

### RNA extraction and QRT-PCR

Extraction of RNA from skin and HSO was done using the Qiagen miRNeasy kit and QIAzol reagent. Samples were disrupted and homogenized in QIAzol buffer. QRT-PCR analysis of target genes was done using a LightCycler96 RT-PCR machine. Primers were designed using NCBI BLAST or publicly available primer sequences. All samples were run in duplicates and equal loading was normalized to expression of housekeeping gene RPL27.

### GTRD database analysis

To analyze the potential role of NF-kB and AP1 transcription factors in the AA skin DEG regulation (DEGs are from our GSE120783 study], we employed GTRDs webtool (http://gtrd.biouml.org/) and selected meta-clusters and maximum gene distance of 5000 nucleotides to define transcription initiation sites in DEGs. We searched for putative binding sites for the human transcription factors RELA (P65), cFOS and cJUN. within −100 - +1000 bases from DEG promoters.

### Statistical Analysis

Data visualization and statistical analysis was conducted using GraphPad PRISM. Western Blot and Microscopy images were analyzed using FIJI (ImageJ). P values were assessed using Welch’s T-Tests. P values <0.05 were considered significant.

## DATA AVAILABILITY STATEMENT

All raw and preprocessed RNASeq data used in this study were previously published and are deposited in the GEO database with the GEO accession number GSE120783. All other data generated or analyzed during this study are included in this article and in appendix files.

## ACKNOWLEDGMENTS

We acknowledge Northwestern University SBDRC (P30AR075049) STEM and TEST IT Cores for technical support. We are thankful to Diya Goyal and Korvell Russell for technical assistance.

The present work is supported by 5R21AR081520, 5R21AI168798, and R01AI125366 (to IB) and UC2AR081033, R01AR080662 and P30AR075049 (to LT).

## DECLARATIONS

### Ethics approval and consent to participate

Informed consent for human biospecimens was obtained in writing following all local and federal regulations under STU00009443 approved by the Northwestern University Institutional Review Board. Neonatal foreskin tissue is approved for collection under STU00009443 with a waiver of consent.

### Author contributions

Conceptualization: IB; Formal Analysis: DT, LCT; Funding Acquisition: IB; Investigation: DT, PG, RK, AK; Methodology: IB, BPW; Validation: DT, PG; Visualization: PG, BPW; Writing - Original Draft Preparation: IB; Writing - Review and Editing: IB, BPW, PG

## Abbreviations

AA: African American
AD: atopic dermatitis
DEG: differentially expressed gene
FC: fold change
GSEA: gene set enrichment analysis
HSO: 3D human skin organoid
IL1R1: Interleukin 1 Receptor Type 1
IL22R1: Interleukin 22 Receptor Subunit Alpha 1
IRAK1: Interleukin 1 Receptor Associated Kinase 1
LCE: late cornified envelope
NHEK: normal human epidermal keratinocytes
qRT-PCR: quantitative real-time reverse-transcription polymerase chain reaction
TF: transcription factor
TNFα: tumor necrosis factor alpha
WNH: White Non-Hispanic

## SUPPLEMENTAL DATA

### Supplemental Materials and Methods

**Supplemental Table 1. Differentially expressed proteins in adult AA skin.** Proteins were extracted from full-thickness skin biopsies from the upper arm skin in AA and WNH healthy female volunteers. Samples were homogenized in a Qiagen Tissue PowerLyzer at 3,500 rpm in TRIzol according to the manufacturer’s protocol (Thermo Fisher Scientific). Total protein was isolated according to manufacturer’s protocol and eluted in 1X PBS. 50µl protein extracts diluted to 1mg/ml/sample were loaded on a 96-well PCR plate and submitted to Olink for quantification of 384 proteins according to their Explore 384 Inflammation panel. Differential protein expression (DEP) analyses were performed using R software (www.R-project.org).

**Supplemental Figure 1. Analysis of NF-kB and AP1 binding sites in DEG promoters using GTRD (Gene Transcription Regulation Database).** Search was done for the putative binding sites within DEG promoters [-1000,+100] for the previously published AA adult skin DEGs [GSE120783] as described in Materials and Methods.

