## Supplemental Table 1 for "Activation of IL1A/IRAK1 axis and downstream proinflammatory signaling in healthy adult and neonatal African American skin"

| Protein/Gene | FC | P Value | Function | Gene expression in tissues | Function in skin | GWAS/genetically associated diseases related to skin | Correlation with skin disease |
|---|---|---|---|---|---|---|---|
| IL1A | 3.98 | 0.037 | Pleiotropic cytokine involved in immune response, inflammation, hematopoiesis. Bridges innate and adaptive immune systems. Role as Alarmin. Can act as a transcription factor. | Ubiquitous expression<br>Higher than average in skin cells including keratinocytes | Inflammation and cell death | Asthma, Psoriasis, Contact Dermatitis, Rheumatoid arthritis | Correlation with psoriasis, AD and Lupus |
| IRAK1<br>Interleukin 1 Receptor Associated Kinase 1 | 2.04 | 0.0009 | Ser/Thr kinase associated with innate immune response via IL1R and Toll-like receptor signaling | Low expression in skin<br>High expression in lymphoid cells | Inflammation | RA , SLE (Lupus) Celiac disease | Increased in psoriatic skin |
| IL4 | 3.25 | 0.036 | Cytokine involved in hematopoiesis inflammation, and T-cell responses | Ubiquitous expression | Drives T-cell differentiation and IgG-e production | Asthma, Psoriasis, Crohn's, Allergy | Involved in AD pathology |
| IL22RA1<br>Interleukin 22 Receptor Subunit a1 | 1.72 | 0.041 | Component subunit of IL20,22,24. IL22 receptor signals through JAK/STAT and ERK1/2 | Ubiquitous expression<br>Higher than average in skin | Promotes keratinocyte proliferation and regulates innate immune responses | Psoriasis, Crohn's, Ulcerative Colitis, Diabetes | Contributes to psoriasis, AD and contact dermatitis |
| AOC1/DAO<br>Diamine Oxidase 1 | 2.73 | 0.053 | Membrane glycoprotein that deaminates histamine and related molecules | Ubiquitous expression | Histamine deamination prevents allergic reactions | Allergic responses | Increased severity of skin symptoms of fibromyalgia (rash, dryness/hyperhidrosis) |
| FST<br>Follistatin | 2.73 | 0.019 | FST negatively regulates the signaling of TGF-β family members: activin, bone morphogenic proteins (BMP), and myostatin | Ubiquitous expression<br>Higher than average in skin (keratinocytes, fibroblasts) | Skin and appendages development, wound healing and inflammation | Acne | |
| YTHDF3<br>RNA binding protein | 3.229 | 0.045 | Binds to m6A in RNA to promote stability by marking RNA for degradation | Ubiquitous expression | Mediates Type 1 Interferon signaling | No specific SNPs identified | Promotes melanoma metastasis through the LOXL3 axis |
| RABGAP1L<br>RAB GTPase Activating Protein 1 Like | 1,49 | 0.018 | Plays a role in endocytosis and intracellular protein transport. Enables continuous directional cell migration via recycling of fibronectin receptors | Ubiquitous expression<br>Very low in skin | | Cholesteatoma | Overexpression in skin cancer (SCC) |
| ICA1<br>Islet Cell Autoantigen 1 | 1.93 | 0.018 | May play a role in neurotransmitter secretion | Pancreas, prostate, testis, brain, very low in skin | ? | Autoantigen in insulin-dependent diabetes mellitus, primary Sjogren's syndrome | ? |
| NELL2<br>Neural EGFL Like 2 | 0.59 | 0.0013 | Affects neuronal cell proliferation, differentiation and survival | Ubiquitous expression | Not specifically related to any skin functions | No specific SNPs identified | Protein marker of AD skin |
| CLSTN2<br>Calsyntenin 2 | 0.27 | 0.039 | Adhesion molecule which promotes synapse formation and calcium ion binding | Ubiquitous expression across many tissues | Not specifically related to any skin functions | Asthma | Not identified in any skin disorders |
