## Supplementary figures and images for "Activation of IL1A/IRAK1 axis and downstream proinflammatory signaling in healthy adult and neonatal African American skin"

### Supplemental Figure 1

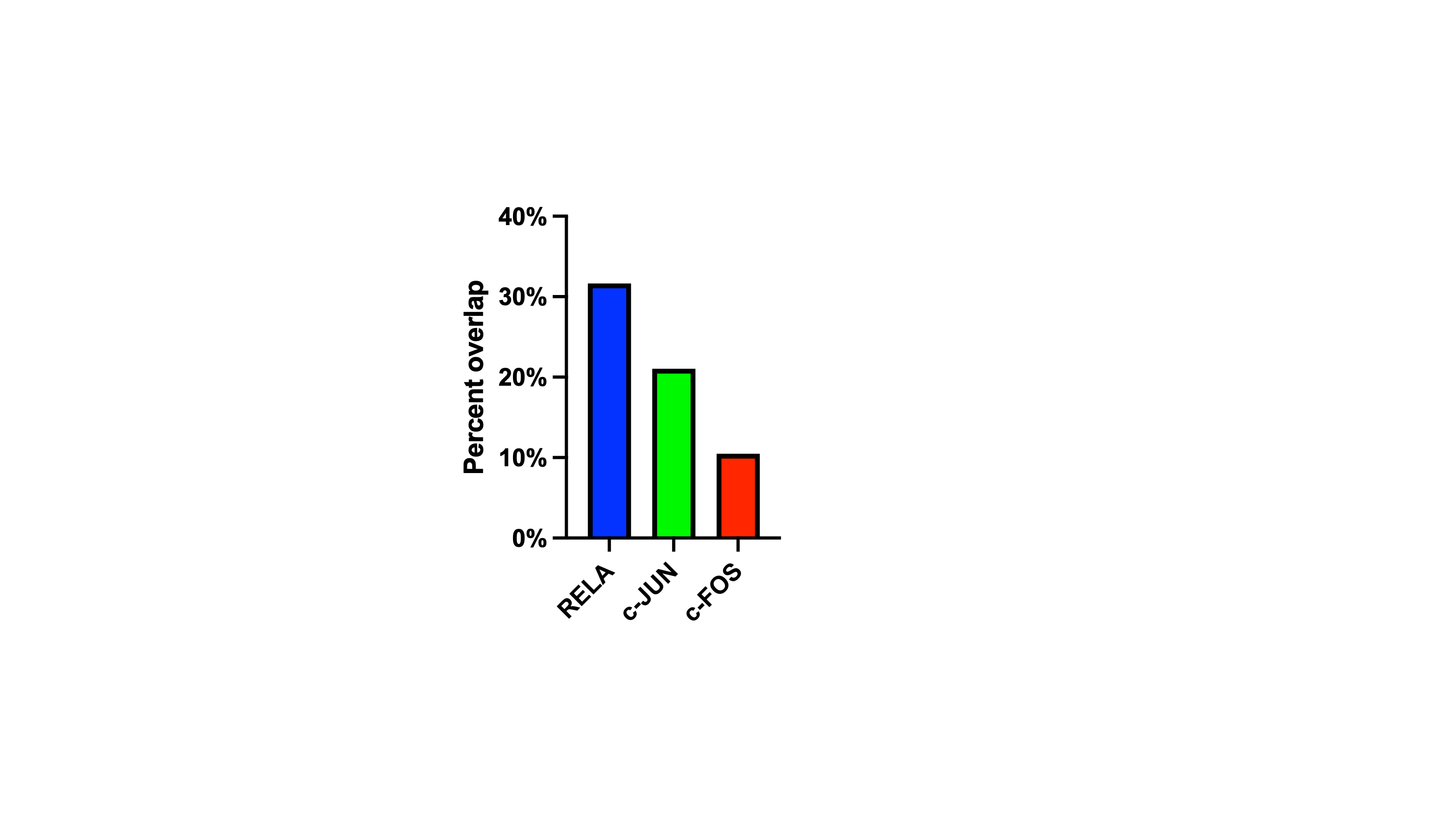
